# Examining *Fouquieria splendens* in an environmental and ecological context: Effect of topography and interspecific neighbors on ocotillo morphology and distribution

**DOI:** 10.1101/2020.11.28.397018

**Authors:** Anastasia Bernat, Acacia Tsz So Tang, Allegra Steenson, Eric Larsen

## Abstract

*Fouquieria splendens* is a stem-succulent native to the Chihuahuan, Mojave, and Sonoran Deserts that spans Mexico and the American Southwest. It is well-known for its variable morphology, the underlying reason for which remains incompletely understood. Here, we attempt to quantify the effect of topographic and interspecific factors on *F. splendens* morphology and distribution. To this end, we measured 27 ocotillos located in the Organ Pipe Cactus National Monument within the Sonoran Desert during June of 2019. We also quantified the spatial distribution of interspecific neighbors relative to *F. splendens* within two topographically different sites: a bajada gradient and a plain. Using ocotillo morphology, the distances to the nearest neighbors of ocotillos, and hydrographic data extracted from the National Hydrography Dataset, we demonstrate 1) the effect of major interspecific neighbors, i.e. shrubs and cacti, on ocotillo morphology; 2) the effect of elevation on intraspecific spacing as individuals compete for limited space; and 3) a trade-off between height and number of branches. This places *F. splendens* morphology in its larger environmental and ecological context, highlighting the importance of individual traits and associated trade-offs among traits affected by topography and interspecific neighbors. By examining the ocotillo in a multi-species community and diverse landscape, this study provides empirical insight into a wider range of factors contributing to the variation in *F. splendens* morphology and spacing.

## Resumen

*Fouquieria splendens* es una planta suculenta nativa de los desiertos de Chihuahua y Sonora que se extienden por México y el suroeste de Estados Unidos. Es reconocida por su morfología variable, que continúa siendo no explicada del todo. Aquí intentamos cuantificar el efecto de factores topográficos y interespecíficas sobre la morfología y distribución de *F. splendens*. Para esto, medimos 27 ocotillos ubicados en el Monumento Nacional Organ Pipe Cactus dentro del Desierto de Sonora durante junio de 2019. También intentamos cuantificar la distribución espacial de los vecinos interespecíficas a *F. splendens* dentro de dos sitios topográficamente diferentes: una pendiente y una llanura. De acuerdo con la morfología del ocotillo, las distancias a los vecinos más cercanos de un ocotillo y los datos de la cuenca hidrográfica, extraídos del Conjunto de Datos Hidrográficos Nacionales, demostramos 1) el efecto de los principales vecinos interespecíficas, como los arbustos y cactus, en la morfología del ocotillo; 2) el efecto de la elevación en el espaciamiento entre vecinos de ocotillo más cercanos a medida que los individuos compiten por un espacio limitado; y 3) una compensación entre la altura y el número de ramas. Esto se ajusta a la morfología de *F. splendens* en un contexto ambiental y ecológico más amplio, lo que ayuda a entender la importancia de los atributos fenotípicos y compensaciones asociadas entre los atributos fenotípicos, afectados por las limitaciones de la topografía y los vecinos interespecíficas. Al examinar el ocotillo en una comunidad multiespecie y paisaje diverso, este estudio presenta una gama más amplia de causas que influyen en la variación en la morfología y el espaciamiento de *F. splendens*.

*Fouquieria splendens* (ocotillo) is a drought-deciduous stem-succulent endemic to the Chihuahuan, Mojave, and Sonoran Deserts and is known for its unusual and highly variable morphology (Killingbeck, 2017). *F. splendens* exhibits stochastic production of segments, inflorescence, and branches (Darrow, 1943; Bowers, 2005; Killingbeck, 2017), which has made the ocotillo statistically and ecologically challenging to study. Past research has focused on the punctuated episodes of terminal segment production in ocotillos to better determine their growth patterns and causal effects (Darrow, 1943; Bowers, 2016; Killingbeck, 2016; Killingbeck, 2019). These episodes of segment growth are marked by ring-like seams which circumscribe the branch and are an attractive feature for dendroclimatic or desert ecology research (Kadmon, 1995; Bowers, 2005). However, conclusions on the effects on ocotillo morphology have been indeterminate, as Killingbeck found that no measurable segment elongation occurred in new or extant branches in some ocotillos for 4-9 years (Killingbeck, 2017). As a result, despite the inclusion of detailed measurements on annual and seasonal rainfall, the variation in the morphology of ocotillos does not appear to be dominated by any singular or seasonal event. Thus, it is important to better understand ocotillo morphology in order to fully determine its ecological function in Southwest deserts and to test general ecological dynamics in multi-species desert communities.

As shown in past studies, the variable growth form of *F. splendens* leads to challenges both in the field and during analysis. For instance, it is not possible to infer annual segment growth measurements because not all branches grow new segments in the same year (Nobel and Zutta, 2004; Killingbeck, 2017). Congruently, the production of new growth and elongation of extant segments show high interplant and intraplant variability (Bowers, 2005). Previous research on ocotillo inflorescence found that variation among branches on a single plant overwhelmed any differences between individuals (Bowers, 2005; Bowers, 2006; Killingbeck, 2009). Studies also indicate that segment growth produced during one year had little or no relation to the amount produced during the succeeding year (Darrow, 1943; Killingbeck, 2017). Furthermore, production of terminal segments increased in stochasticity with maturity, and growth ceased after branches attained a length of 3-4.5 m (Darrow, 1943). In turn, the reasons behind this variability, whether genetic or environmental, remain inconclusive, and it is in our benefit to further explore which additional factors could be influencing or directing ocotillo growth.

Studying ocotillo growth in the Sonoran Desert is favorable because it is a multi-species desert with long mountain ranges and low-elevation valleys (Bowers and McLaughlin, 1982). Its topography makes it ideal to study ocotillo morphology near arroyos or along bajadas. Arroyos are characterized as intermittently dry creek beds, and bajadas are slopes that exhibit a gradient of soil particle size and water availability (Phillips and MacMahon, 1978; McAuliffe, 1994; Stromberg, 2007). These geological features have allowed researchers to study both diverse competition dynamics and topographic effects on the growth of various desert plant species in the field (Bowers and Lowe, 1986; Nobel and Zutta, 2007; Pierce, 2019); however, these features have been overlooked in ocotillo studies.

Wider studies on desert plant communities have investigated the interaction between morphology and competition. In particular, competition in desert communities is a likely candidate to influence plant form and spatial distribution. It has been assumed that deserts are unproductive and exhibit low competition because of their scarcity of water (Grime, 1973; Huston, 1979; Gaudet and Keddy, 1989). However, studies have revealed a diverse spatial distribution of desert plants and have shown how competition among plants can vary in levels of intensity (Fonteyn and Mahall, 1978; Fowler, 1986; Kadmon, 1995; Ji et al., 2019). Plant removal experiments in arid regions have suggested that interspecific competition is more intense than intraspecific competition (Fonteyn and Mahall, 1978) while experiments along productivity gradients revealed a positive relation between competition and plant density (Kadmon, 1995). Furthermore, Pielou’s model provides evidence of a positive correlation between nearest-neighbor distance and plant size (Pielou, 1962), and Philips and MacMahon found that plant size increased as dispersion patterns became more regular (Philips and MacMahon, 1981). Others have also defined specific, competition type interactions; Ji and colleagues (2019) used remote sensing technology to determine that shrub-shrub competition is most likely only active in locations where conditions have highly favored shrub establishment and survival. In turn, to a large extent, plant competition also drives patterns in plant density and size.

Measuring interspecies interactions can help elucidate how desert plants survive or out-compete their potential competitors. Plant fitness has been well-studied and has led to recent novel observations regarding trade-offs between environmental context, competitor defence, plant growth, and plant reproduction on the population level (Lundgren and Marais, 2020; Pearse et al., 2020). All plants must balance the allocation of resources for growth, survival, and reproduction in order to maximize fitness within the constraints of competition. Ge and colleagues extend that concept to a three decade study of trait and competition relationships across 51 winter annual desert species (Ge et al., 2019). They found that trait trade-offs scaled to the community level and were exhibited most by the dominant species (Ge et al., 2019). In turn, trait trade-offs can serve as an organizing principle for understanding plant population and community processes. Within the multi-species communities of the Sonoran Desert, we hypothesized that our empirical approach to studying the ocotillo within its environmental context would help elucidate potential trait trade-offs.

This study examined how environmental and ecological factors such as elevation, intraspecific distances, and interspecific distances affected ocotillo morphology in order to better understand patterns in ocotillo growth form and its larger contribution to the spacing of plants in complex desert plant communities. We measured the height, circumference, number of branches, and terminal segment lengths of ocotillos as well as the nearest intraspecific and interspecific neighbor distances; furthermore, we measured elevation and cross-referenced distances to the nearest arroyo (United States Geological Survey National Hydrography Dataset). Our study focused on major and minor nearby interspecific plants as well as the presence of shrubs vs. cacti. By assessing ocotillo plant form as well as the spatial proximity of ocotillos to their interspecific neighbors across various elevations and sites, we tested how interspecific and topographic differences relate to and effect ocotillo growth form. Here we provide empirical insight on how to examine the ocotillo in its larger environmental and ecological context, which has helped uncover novel patterns in ocotillo morphology and distribution. Results suggest that interspecific factors and topographic differences affect ocotillo morphology and spatial density, and they provide an important framework to reexamine this unique desert plant.

## Materials and Methods

### Study Area

The study area was located in the Sonoran Desert at Organ Pipe Cactus National Monument (31.94515°N, 112.81252°W), approximately 16 km north of the US-Mexico border in southern Arizona (Fig. 1). The plant community contains the most striking diversity out of the four main deserts in the American Southwest (Bowers and McLaughlin, 1982). In addition to *F. splendens*, we observed adjacent, interspecific species. These included *Carnegiea gigantea* (saguaro cactus), *Larrea tridentate* (creosote bush), *Stenocereus thurberi* (organ pipe cactus), *Ambrosia deltoidea* (triangle leaf bursage), *Cylindropuntia fulgida* (jumping cholla), *Jatropha cuneata* (limber bush), *Simmondsia chinensis* (jojoba), *Parkinsonia aculeate* (Mexican palo verde), and *Cylindropuntia leptocaulis* (pencil cholla).

**Fig 1.**
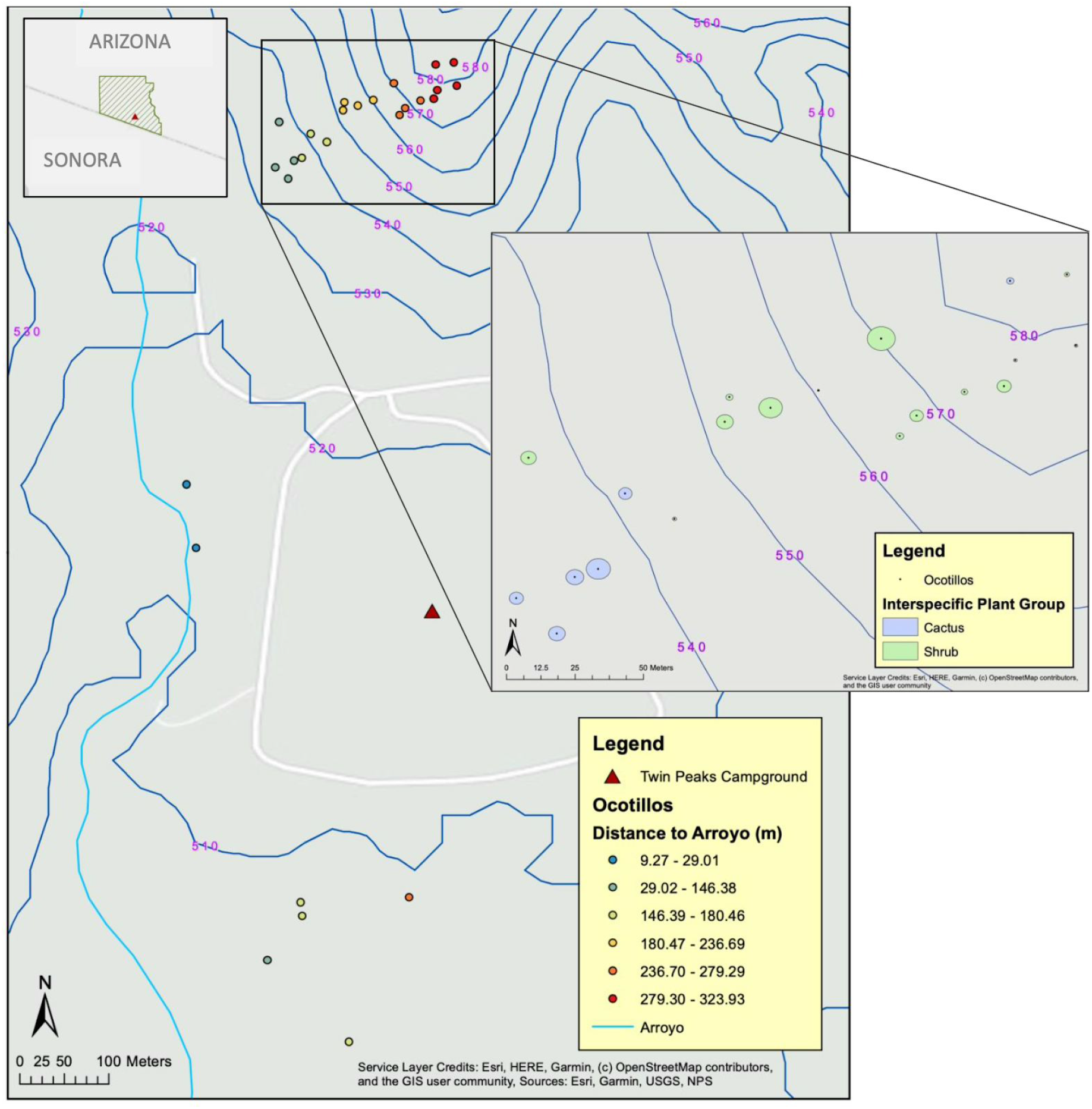
Organ Pipe National Monument Site Map. Top left corner shows the outline of the Organ Pipe National Monument. Waypoints for ocotillos on the bajada, arroyo, and plain are represented by circular points. Bounding box on the right zooms into the ocotillos on the bajada. Hydrographic data were retrieved from the National Hydrography Dataset, symbolized by the blue line, and the topographic basemap was imported from ESRI. The distance from each ocotillo to the arroyo is represented by a color ramp from blue to red corresponding to increasing distance. Additional details such as campground and trail location are included for reference.

We collected data during June 2019 across two topographically distinct sites: the upper section of a bajada (532-581 m) and the lower section of the bajada consisting of a relatively flat plain (501-506 m). In this study, we respectively termed the upper and lower bajada as ‘bajada’ and ‘plain’.

### Data Collection

At each site, we selected ocotillos for measurement and marked their locations using a Garmin GPSMAP 64st. We counted the total number of branches and used a tape measure to take a series of morphological measurements to the nearest cm: height of the ocotillo, circumference at breast height (approximately 1.5 m from the ground), and segment lengths of 10 branches selected uniformly around the ocotillo as illustrated in Figure 2. We quantified interspecific spacing by measuring the distance from an ocotillo to the nearest major interspecific plant and recording the identity of the neighboring plant. We defined a major interspecific neighbor as the tallest and largest-bodied interspecific individual between 1 m and 10 m from the observed ocotillo. We hypothesized that these plants dominated the resources of the local area and could be a source of competition to other flora. Additionally, we identified small interspecific plants up to 30 cm tall within a 1 m radius of the ocotillo as minor interspecific neighbors, which were considered to have a small effect on resources of the local area.

**Fig 2.**
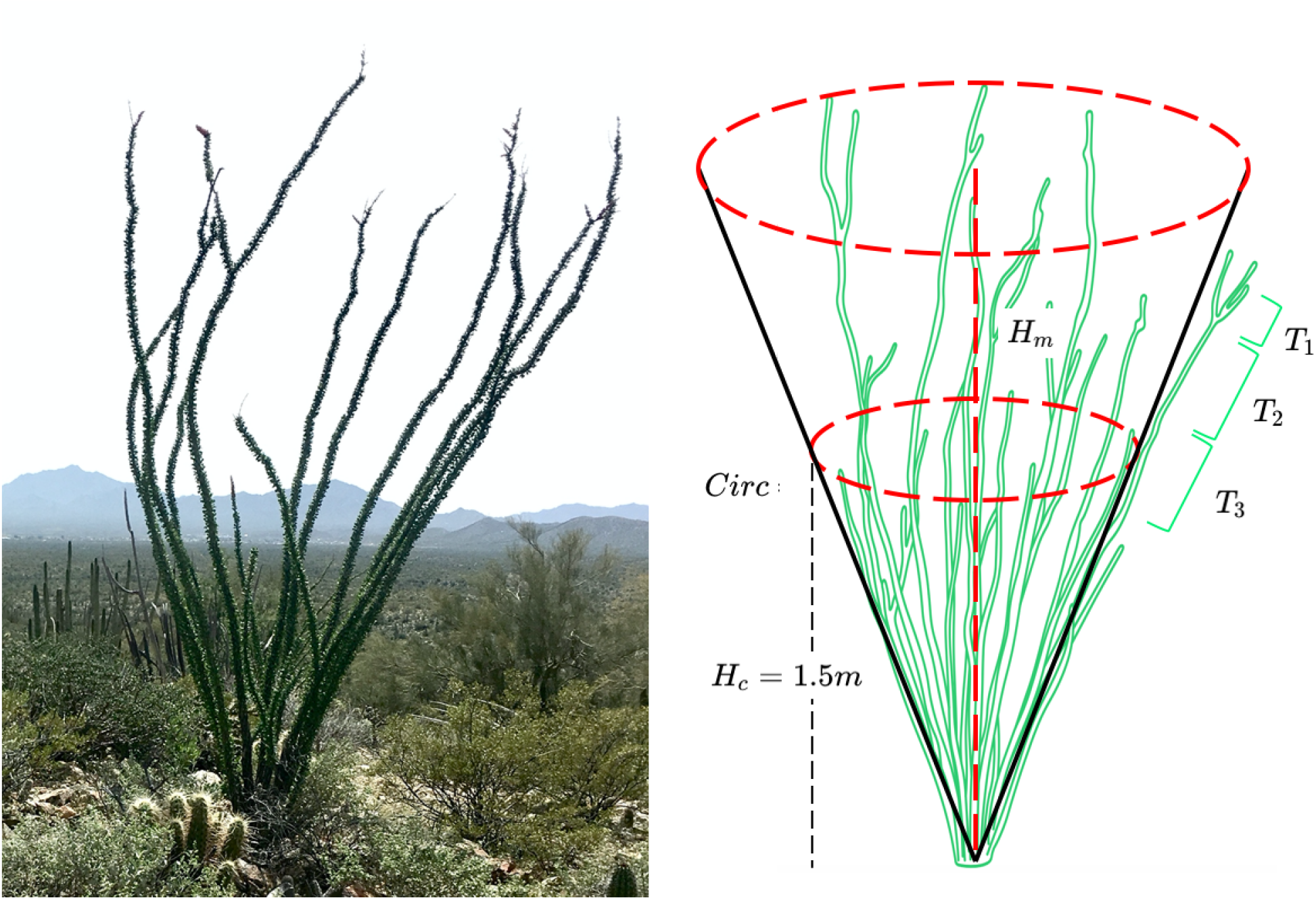
*Fouquieria splendens* morphology measurements. Five types of measurements were taken in the field per ocotillo using measuring tape: 1) circumference (*Circ)* taken at breast height (*H*_*c*_ *=* 1.5 m), 2) the full height of the ocotillo (*H*_*m*_), 3) number of branches, 4) number of segments per 10 branches chosen above breast height, and 5) each of the 10 branches’ segment lengths (*T*_*n*_ where *n* = 1, 2, 3, …).

We measured 20 ocotillos on the bajada and 7 ocotillos on the plain. We traveled down from the peak of a south-facing bajada, selecting an ocotillo for measurement along an elevation contour every 10 m for a total of twenty ocotillos. We measured ocotillos on the bajada over the course of three days. Over the following two days, we measured seven ocotillos on a flat plain at an elevation similar to the bottom of the bajada gradient; two of those seven measured were within 10 m of an arroyo.

### Hydrographic Data

We accessed hydrographic data using the USDA Geospatial Data Gateway and collected from the National Hydrography Dataset (NDH). We cross-referenced the data from the NDH with our ocotillo data in ArcMap 10.7.1 (Esri, 1969) in order to determine the distance from each plant to the nearest arroyo.

### Assumptions

Our data collection methods were based on several assumptions. Plant age was unknown; thus, at best, we can only use proxies of age such as the number of branches on an ocotillo (Darrow 1943). Past research defined mature ocotillos as individuals that are 2.5-4.5 m in height and have 40-75 straight, slender branches (Darrow 1943). Based on that definition, none of our ocotillos would have been mature. In addition, we did not know the year in which each terminal segment was grown. Darrow found low correlation values for branches active in both consecutively-paired years, indicating that the segment growth during one year had little or no relation to the segment produced during the succeeding year (Darrow 1943). Thus, we assumed that segments were not necessarily produced in sequential years.

### Morphology, Proximity, and Topography Analyses

We analyzed the relationships between ocotillos’ morphology, site, elevation, the presence of major shrubs or cacti, proximity to the local arroyo, proximity to other ocotillos, and proximity to interspecific plants; we conducted these analyses using multivariate linear regressions in the program RStudio version 1.2.5033 (RStudio, 2009). Additionally, we calculated for each ocotillo its number of nodes, median branch length, branch length interquartile range (IQR), and terminal segment length IQR; all these calculations were included in the analyses. A node is defined as the ring-like seam that circumscribes a branch. Branch length IQR and terminal segment length IQR represent morphology variation where the larger the IQRs, the more variation in branch length and terminal segment length the ocotillo has, respectively.

We also ran two-sample *t*-tests for categorical variables, i.e. site or nearby major interspecific plant group. All measurements followed log-normal distributions except for circumference, number of nodes, median branch length, and distance to the nearest arroyo, which were normally distributed. After log-transforming all response variables and centering all predictor variables, models constructed passed linear regression assumptions. Selection for the best fit model was based on Akaike information criterion (AIC) and an analysis of variation (ANOVA) test where p > 0.05 favored the simpler model (Supplementary Files).

### Terminal Segment Length Analyses

Terminal segments refer to the last 5 segments on the ends of each of the 10 branches measured per ocotillo; they reflect the most recent growth in the plants, and we observed them to be the most distinct and easy to measure. We analyzed the terminal segment lengths using mixed-effect modeling to determine the effects of elevation, site, ocotillo morphology, and interspecific plant type as well as distance to the nearest arroyo, ocotillo neighbor, or interspecific neighbor (Supplementary Files). To account for pseudo-replication in the models, we designated ocotillo number, segment number, and branch number as random effects. To further examine interspecific impact, we tested the effect of major interspecific plant type - either a shrub or cactus - on ocotillo terminal segment length. This was done using a Kruskal-Wallis test because the data are non-normally distributed (Shapiro-Wilk’s normality test p<0.05).

### Principal Components Analysis

We carried out two principal components analyses (PCAs) to describe the morphology of the individuals measured (Supplementary Files). The first included six variables: height, circumference, number of branches, median terminal segment length, median branch length, and number of nodes. The second included the aforementioned variables and two additional variables: branch length IQR and terminal segment length IQR. We scaled data such that each variable had the same variance (unit variance). This prevented variables with larger variances to dominate the PCA, and it was necessary because variables, such as height and segment length, had different magnitudes. After the analysis, we defined principal components (PCs) that accounted for sufficient variation as those with eigenvalues larger or equal than 1, meaning that such PCs capture more variation in the data than the original variables (Kaiser, 1960; Kaufman and Dunlap, 2000).

## Results

### Site And Median Branch Length Affect Height

The best fit model for height included site and terminal segment length IQR (*n* = 27; *P* < 0.05); however, the terminal segment length IQR of the first ocotillo measured was an outlier (Dixon’s test; *P* < 0.05). We removed this outlier, and found the best fit model for ocotillos across the bajada and plain (*n* = 26) to include site and median branch length (*P* < 0.05). Ocotillos with larger median branch lengths were taller (*β* = 0.002, *SE* = 0.0007, *t* = 2.98, *P* = 0.007; Fig. 3a) where for every 1 m growth in branch length, height increased by 25%. Additionally, ocotillos growing on the plain (*n* = 7) reached greater heights on average than those growing on the bajada (*mean*_plain_ = 1.37 m; *mean*_bajada_ = 1.03 m; *t*(12) = −4.16, *P* = 0.001; Fig. 3b). Ocotillo height increased by 27% when the site changed from the bajada to the plain (*β* = 0.24, *SE* = 0.08, *t* = 3.16, *P* = 0.004). For ocotillos only located along the bajada (*n* = 20), the best fit model included only median branch length (*β* = 0.002, *SE* = 0.0009, *t* = 2.86, *P* = 0.01).

**Fig 3.**
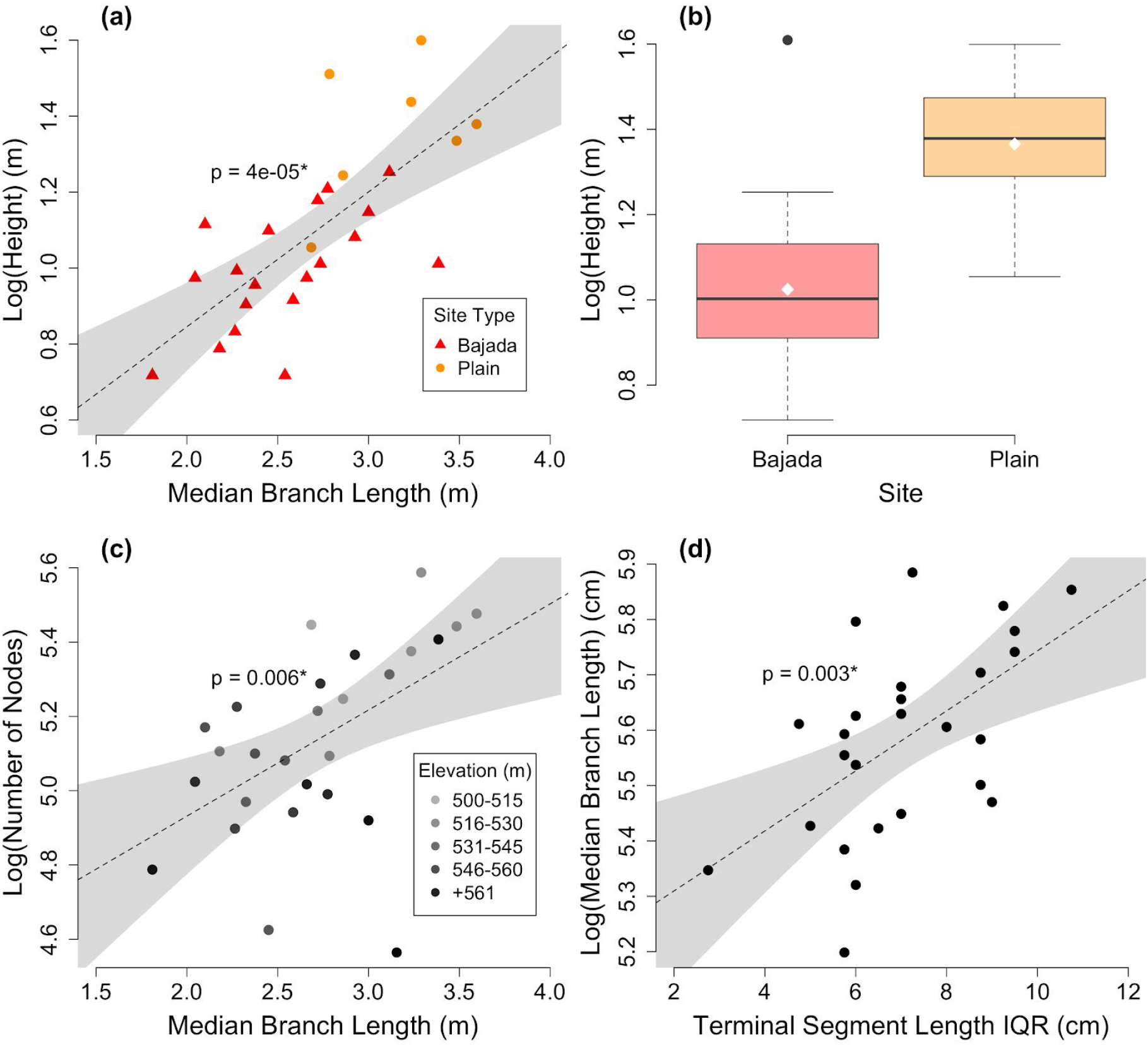
Ocotillo height related by site and median branch length. Data points are coded by landform types where orange circles represent ocotillos on the plain and red triangles represent ocotillos on the bajada. The dashed linear regression lines indicate a significant association (*P* < 0.05); the shaded bands indicate 95% confidence intervals. (a) Median branch length is positively related to height (*n* = 26, *β* = 0.36, *SE* = 0.07, *t* = 5.0), but this is largely due to differences in site. (b) There was a significant effect of site where ocotillos on the plain exhibited higher heights on average compared to ocotillos on the bajada (*mean*_plain_ = 1.37 m; *mean*_bajada_ = 1.03 m; *t*(12) = −4.16, *P* = 0.001). The box represents the 25th and 75th percentiles, the whiskers the 10th and 90th percentiles, dots the 5th and 95th percentiles, the white diamond is the mean, and the solid line is the median. (c) Median branch length is positively related to the number of nodes (*n* = 27, *β* = 0.29, *SE* = 0.09, *t* = 3.0). (d) Terminal segment length interquartile range is positively related to median branch length (*n* = 27, *β* = 0.36, *SE* = 0.02, *t* = 3.4).

### Branch Length and Elevation Affect Number of Nodes

For ocotillos along the bajada and plain, the best fit model for the number of nodes included elevation and median branch length (*n* = 27; *P* < 0.05). For every 10 m increase in elevation, the number of nodes on an ocotillo decreased by 4% (*β* = −0.004, *SE* = 0.002, *t* = −2.75, *P* = 0.01). Additionally, for every 1 m growth in median branch length, the number of nodes increased by 23% (*β* = −0.002, *SE* = 9e-04, *t* = 2.39, *P* = 0.03; Fig. 3c). For ocotillos only along the bajada, the best fit model included median branch length and branch length IQR (*n* = 19; *P* < 0.05). Similarly, for every 1 m growth in median branch length, the number of nodes increased by 34%, but the effect was stronger (*β* = −0.004, *SE* = 0.002, *t* = −2.75, *P* = 0.01). Conversely, for every 1 m increase in branch length IQR, the number of nodes decreased by 28%, suggesting an increase in the variation in branch length leads to a decrease in the number of nodes.

### Height, Number of Nodes, and Terminal Segment Length IQR Affect Median Branch Length

For ocotillos across the bajada and plain, the best fit model for median branch length included height, number of nodes, and terminal segment length IQR, which were all positively related to median branch length (*n* = 26; *P* < 0.05). Across all measurements, median branch length was the only measurement strongly related to terminal segment length IQR (Fig. 3d). For every 10 cm growth in terminal segment length IQR, the ocotillo increased its median branch length by 51% (*β* = 0.04, *SE* = 0.01, *t* = 4.05, *P* = 5.3e-04), suggesting that an increase in recent segment length variation leads to longer branch lengths overall. Additionally, similar aforementioned relationships were reflected in the model. For every 1 m growth in height, the median branch length (cm) increased by 7.3% (*β* = 0.07, *SE* = 0.03, *t* = 2.30, *P* = 0.03) and for every 20 nodes grown, the median branch length (cm) increased by approximately 4% (*β* = 0.002, *SE* = 5.4e-04, *t* = 3.75, *P* = 0.001). Likewise, for ocotillos across the bajada, the best fit model of median branch length included the number of nodes and terminal segment length IQR (*n* =19, *P* < 0.05), but their effects were stronger. For every 10 cm growth in terminal segment length IQR, median branch length grew by 63% (*β* = 0.05, *SE* = 0.01, *t* = 3.57, *P* = 0.003); additionally, for every 20 nodes grown, the median branch length (cm) increased by 5.2% (*β* = 0.003, *SE* = 8e-04, *t* = 3.27, *P* = 0.005)

### Number of Nodes, Branch Length IQR, and Number of Cacti Affect Terminal Segment Lengths

The best fit, mixed-effect model of terminal segment length for all ocotillos except the first ocotillo measured (*n* = 1,300) included the number of nodes, branch length IQR, and the number of cacti within 1 m from the base of the ocotillo. Random effects were ocotillo number and segment number. The minor, interspecific measurement was not significant as a sole predictor, but it was as an interaction term with the number of nodes (*β* = 0.05, *SE* = 0.01, *t* = 3.76). For every additional cactus around the base of the ocotillo and 20 new nodes grown, the terminal segment length increased by 1 cm. However, as a sole predictor, the number of nodes was negatively related to terminal segment length (*β* = −0.06, *SE* = 0.01, *t* = −4.81) where for every 20 new nodes grown, the terminal segment length decreased by 1.2 cm. Finally, each additional 1 m growth in branch length IQR increased ocotillo terminal segment length by 3.6 cm (*β* = 0.04, *SE* = 0.01, *t* = 2.57).

For ocotillos on the bajada, the best fit model of terminal segment length included the number of nodes and the number of minor cacti (*n* = 950; *P* < 0.05) with the random effects of ocotillo number and segment number. This model received two times more support than the second-ranked model, which included the number of nodes and branch length IQR. On the bajada, the number of cacti were instead negatively related to terminal segment length (*β* = −1.2, *SE* = 0.57, *t* = −2.07) where for approximately every 1 additional cactus around the base, the terminal segment length decreased by 1 cm. Similar for all ocotillos, for every 20 new nodes grown, the terminal segment length decreased by 1.3 cm.

From Kruskal-Wallis tests, we found ocotillos whose interspecific neighbor was a shrub had more extreme terminal segment lengths than ocotillos whose interspecific neighbor was a cactus (*n* = 1,349, *P* = 0.023; Fig. 4). There was also a significant difference for terminal segment lengths between ocotillos that grew on the bajada or plain (*n* = 1,349, *P* = 0.002) where ocotillos on the bajada reached more extreme terminal segment lengths.

**Fig 4.**
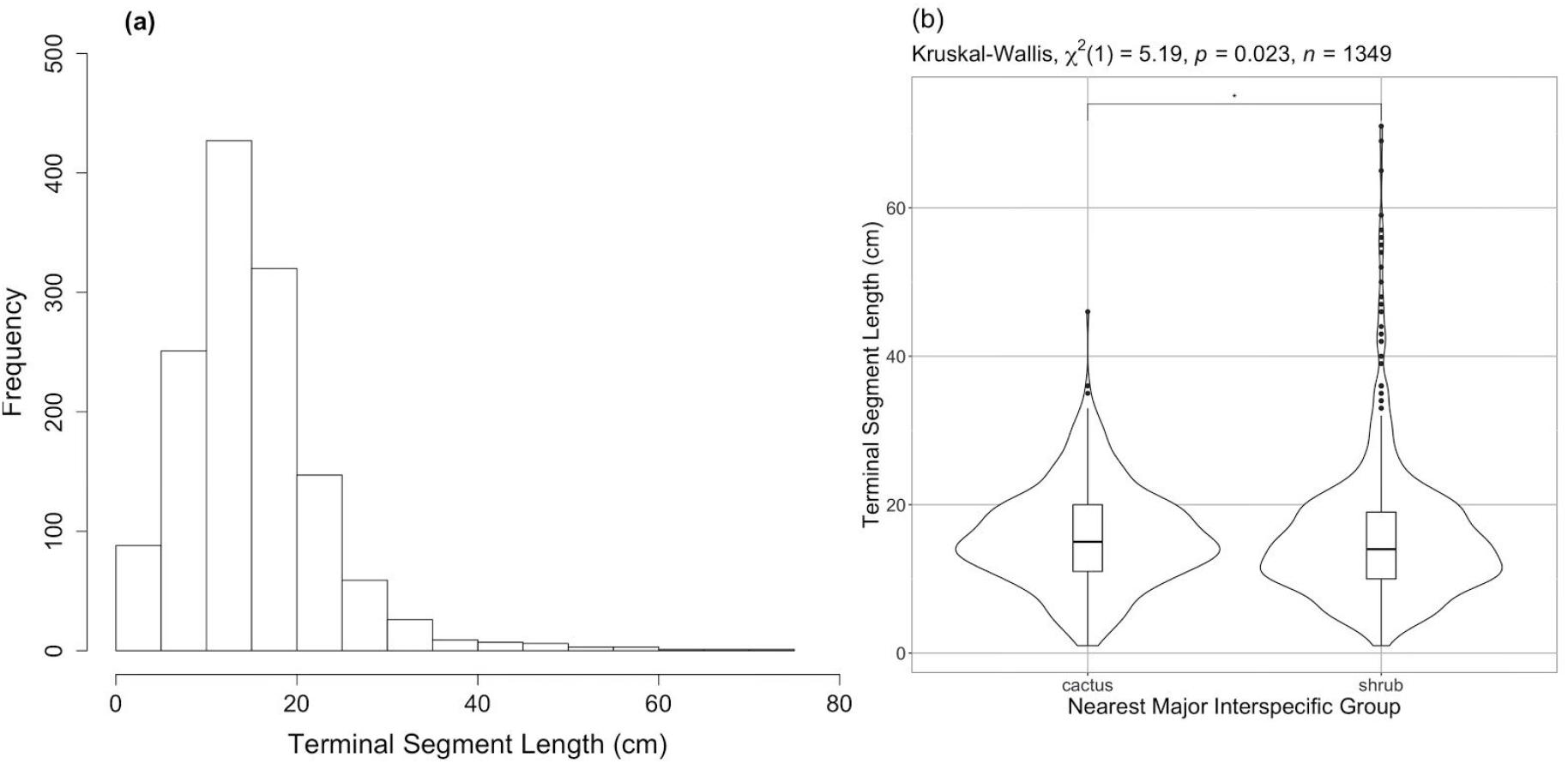
Terminal segment length skewness and nonparametric interspecific plant group effects. (a) Histogram of terminal length showing a long right tail. (b) Violin plot of terminal segment lengths among ocotillos whose nearest, major, interspecific neighbor was either a cactus or shrub.

### Number of Branches Affects Circumference

For ocotillos across the bajada and plain (*n* = 27), the best fit model for circumference included only branch number. Ocotillos with more branches had larger circumferences (*β* = 0.015, *SE* = 0.005, *t* = 2.88, *P* = 0.008; Fig. 5), and for every branch an ocotillo grows, the circumference (m) of the ocotillo grows by 1.54%. The same best fit model also best explained bajada ocotillos (*n* = 20). All other measurements were unrelated to circumference for all ocotillos and for ocotillos on the bajada.

**Fig 5.**
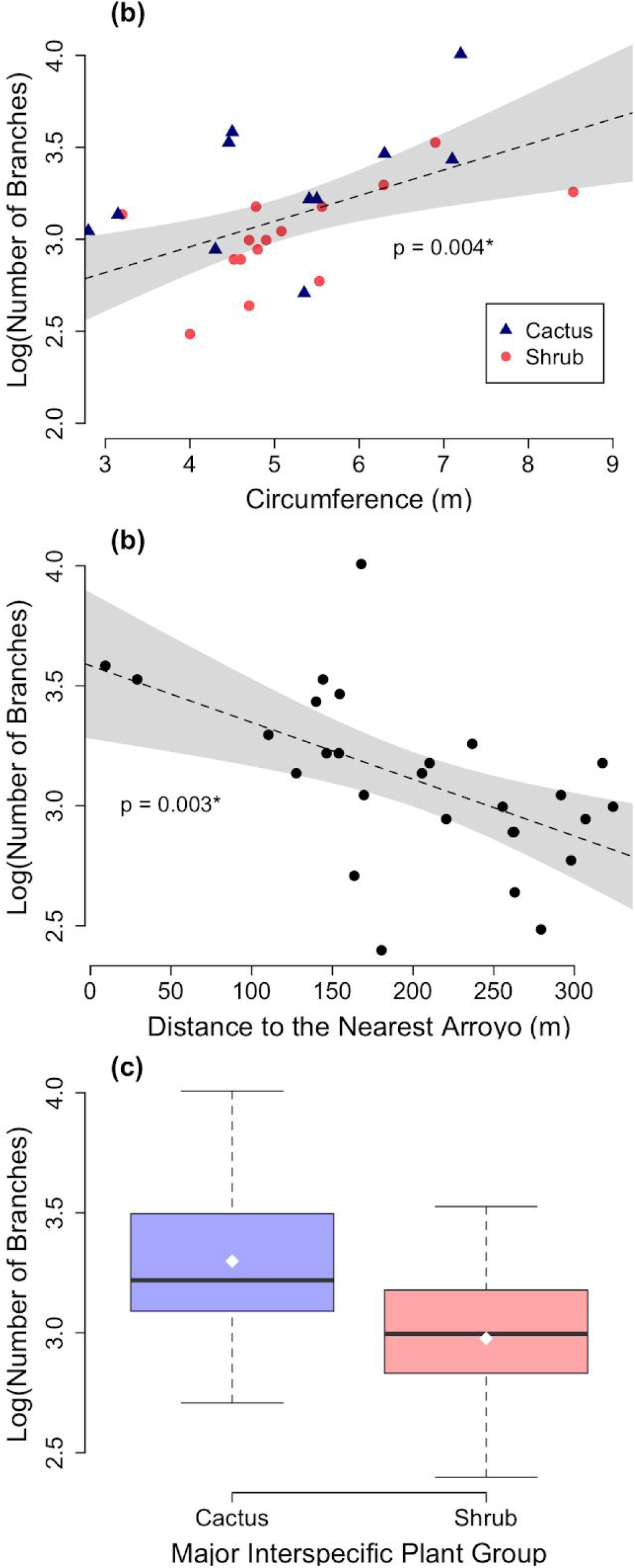
Number of branches related by circumference, distance to the nearest arroyo, and interspecific plant group (e.g. shrub or cactus). The dashed linear regression lines indicate a significant association (*P* < 0.05) and the shaded bands indicate 95% confidence intervals. (a) Circumference is positively related to branch number (*β* = 0.14, *SE* = 0.04, *t* = 3.21). Data points are also coded by interspecific plant groups where circles represent ocotillos with major interspecific neighbors that are shrubs and triangles represent ocotillos with major interspecific neighbors that are cacti. (b) Distance to the nearest arroyo is negatively related to branch number (*β* = −0.003, *SE* = 0.0007, *t* = −3.30).(c) There was a significant effect of interspecific plant group where ocotillos with a cactus exhibited more branches on average compared to ocotillos with a shrub (*mean*_cactus_ = 3.30; *mean*_shrub_ = 2.98 *t*(19.1) = 2.47, *P* = 0.02). The box represents the 25th and 75th percentiles, the whiskers the 10th and 90th percentiles, dots the 5th and 95th percentiles, the white diamond is the mean, and the solid line is the median.

### Circumference, Major Interspecific Group, and Arroyo Distance Affect Number of Branches

For ocotillos along the bajada and plain (*n* = 27), the best fit model for the number of branches among ocotillos included interspecific distance, distance to the nearest arroyo, and the circumference of the ocotillo. This model received over two times more support than the second-ranked model, which included only circumference and arroyo distance. Individual ocotillos with a wider circumference (m) had more branches (*β* = 0.14, *SE* = 0.045, *t* = 68.57, *P* = 3.76e-04). In addition, for every 10 m an ocotillo grew away from its nearest arroyo, the number of branches decreased by 1.63% (Fig. 5). Major interspecific distance was only marginally significant in the model (*β* = 0.067, *SE* = 0.036, *t* = 1.84, *P* = 0.079). Furthermore, among ocotillos whose major interspecific neighbor was a shrub, the average number of branches was lower than ocotillos whose interspecific neighbor was a cactus (*n*_cactus_ = 11; *n*_shrub_ = 16; *mean*_cactus_ = 3.30; *mean*_shrub_ = 2.98; *t*(19.1) = 2.5, *P* = 0.02, Fig. 5c).

For ocotillos on the bajada (*n* = 20), the best fit model included the interspecific plant group as a sole predictor and the interaction term between circumference and the distance to the nearest neighboring ocotillo. This model received twice more the support as the second-ranked model, which included elevation. Within the best fit model, there was a significant positive effect of circumference (*β* = 0.12, *SE* = 0.037, *t* = 3.27, *P* = 0.005) where for every 1 m increase in circumference, the number of branches increased by 12.7%. There was also a weak, negative, interaction effect between circumference and the distance to nearest ocotillo (*β* = −0.02, *SE* = 0.01, *t* = −2.25, *P* = 0.04) where ocotillos with larger circumferences that aggregated closer together exhibited fewer branches. No other effects were significant.

### Elevation and Ocotillo Size Relates to Nearest Ocotillo Distance

For ocotillos along the bajada and on the plain (*n* = 27), the best fit model of the distance to the nearest neighboring ocotillo included the height and circumference of the ocotillo as well as elevation. Individual ocotillos with wider circumferences and taller heights were farther away from neighboring ocotillos (*β* = 0.56, *SE* = 0.19, *t* = 2.99, *P* = 0.0069) where for every 1 m increase in circumference and height, the nearest neighboring ocotillo distance increased by an estimated 75%. Ocotillos at higher elevations and with wider circumferences were also farther from neighboring ocotillos, but the relationship was marginally significant (*β* = 0.011, *SE* = 0.005, *t* = 2.03, *P* = 0.056).

For ocotillos along the bajada (*n* = 20), the best fit model for distance to the nearest neighboring ocotillo included branch number, elevation and distance to the nearest, major interspecific neighbor. Ocotillos higher up in elevation had closer neighboring ocotillos (*β* = −0.02, *SE* = 0.007, *t* = −2.66, *P* = 0.018; Fig. 6a). For every 10 m increase in elevation, the nearest neighboring ocotillo decreased its distance by an estimated 18%. Individuals with more branches and with farther interspecific neighbors also had closer neighboring ocotillos (*β* = −0.062, *SE* = 0.019, *t* = −3.26, *P* = 0.005; Fig. 6b). All remaining predictors in the best fit model were not significant.

**Fig 6.**
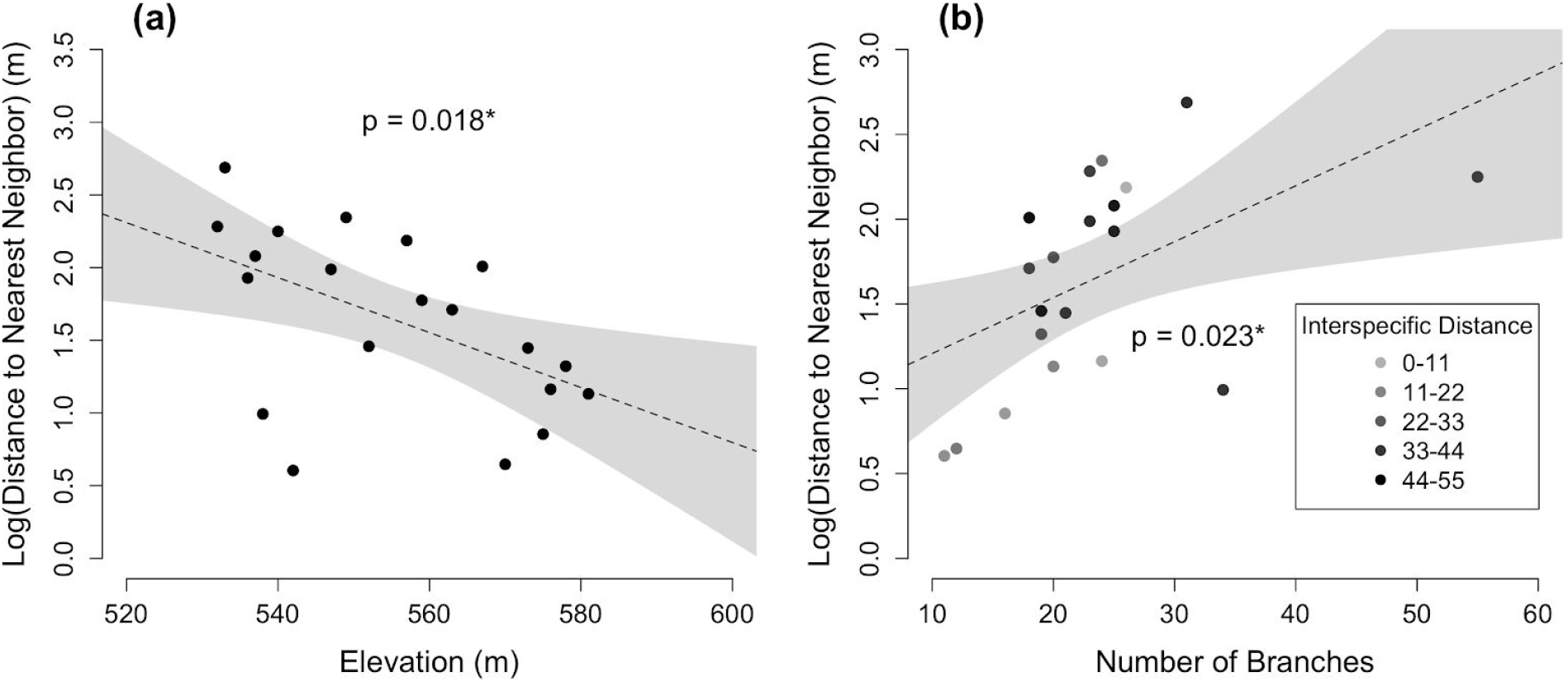
Regressions of distance to the nearest neighboring ocotillo with (a) elevation (*n* = 20, *β* = −0.02, *SE* = 0.007, *t* = −2.59) and (b) number of branches (*n* = 20, *β* = 0.033, *SE* = 0.013, *t* = 2.49) for only ocotillos measured on the bajada. The dashed linear regression lines indicate a significant association (*P* < 0.05); the shaded bands indicate 95% confidence intervals.

### Arroyo Distance Relates to Interspecific Distance

The best fit model of distance to the nearest interspecific neighbor among ocotillos includes only the distance to the nearest arroyo. This model received approximately twice as much support as the second-ranked model, which included elevation. Major interspecific neighbors that were close to an arroyo was farther from the ocotillo (*β* = −0.0038, *SE* = 0.0019, *t* = −2.02, *P* = 0.054), but this was marginally significant. However, among bajada ocotillos, the best fit model is the same as with all ocotillos, but the effect is significant and slightly stronger (*β* = −0.0054, *SE* = 0.0023, *t* = −2.36, *P* = 0.030) where for every 20 m closer the interspecific plant is to an arroyo, its distance from the ocotillo increases by 10.25%.

### Principal Component Analysis

For the first PCA (Fig. 7a), we generated six PCs in total. Of these, three (PC1, PC2 and PC3) had eigenvalues above one (Kaiser, 1960; Kaufman and Dunlap, 2000). Together, they capture 84.5% of all variation between individuals. PC1 captures 40.8% of variation within individuals. Main contributions to PC1 are number of nodes (33.8%), median branch length (30.9%), and height (27.4%). It is worth noting, however, the median terminal segment length only has a low cos2 of 0.16 with respect to PC1, indicating a low ability of PC1 to predict variation in median branch length (Abdi and Williams, 2010). PC2 captures 25.3% of variation, and contributions are mainly made by circumference (51.5%) and number of branches (47.4%). PC3 captures 18.3% of the variation. Median terminal segment length (71.6%) and height (13.2%) are the main contributions to PC3.

**Fig 7.**
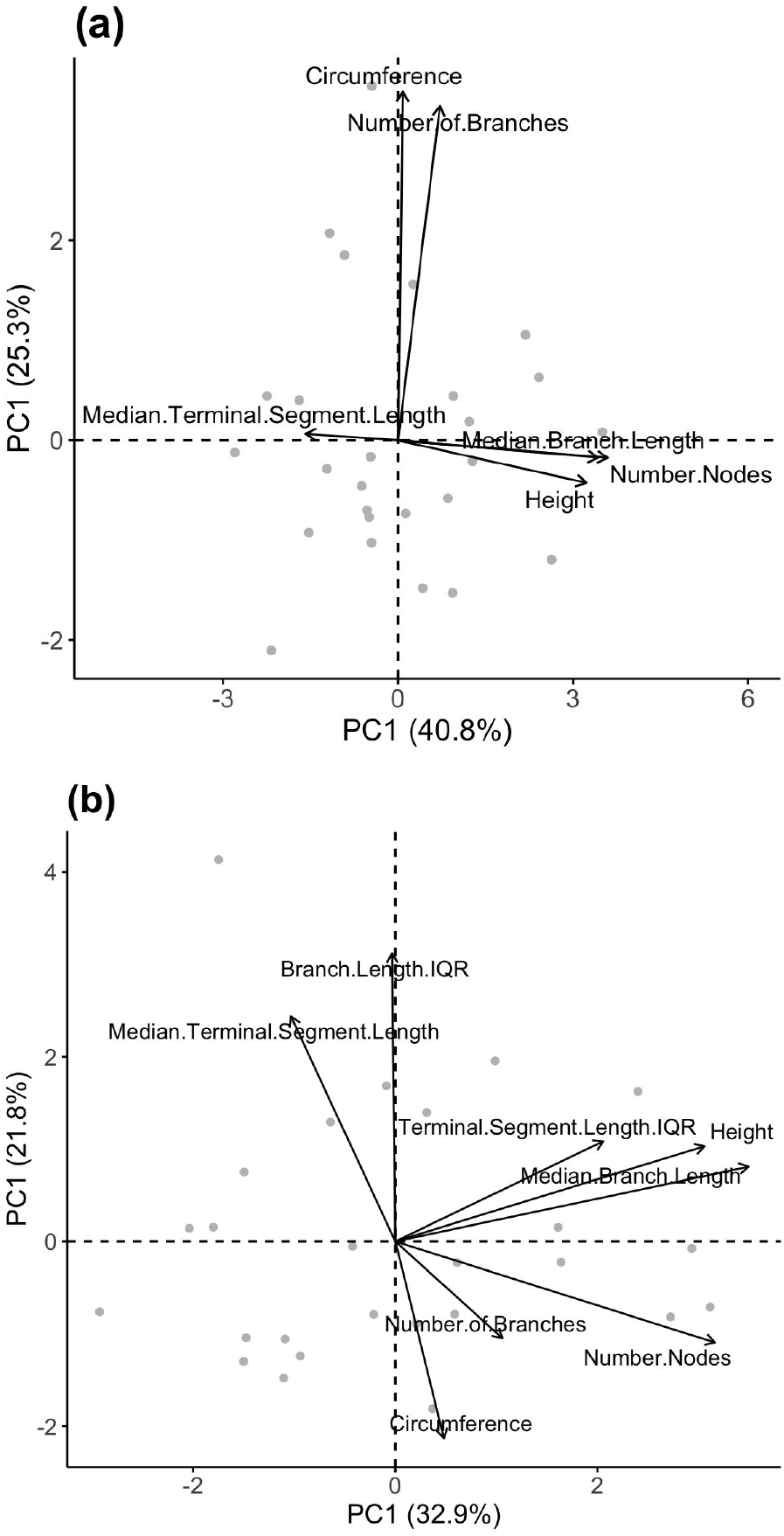
A 2D Principal component analysis (PCA) plot of *F. splendens* morphology. Each dot represents an individual ocotillo. (a) Principal component 1 (PC1) accounts for 40.8% of all variation and is mainly contributed to by number of nodes, median branch length, and height. Principal component 2 (PC2) accounts for 25.3% of all variation and is mainly contributed to by circumference and number of branches. (b) PC1 accounts for 32.9% of all variation and is mainly contributed to by median branch length, number of nodes, height, and terminal segment length IQR. PC2 accounts for 21.8% of all variation and is mainly contributed to by branch length IQR, median terminal segment length, and circumference.

For the second PCA (Fig. 7b), eight PCs were generated. Three of these (PC1, PC2 and PC3) had eigenvalues above one, and together captured 75.4% of all variation. PC1 captures 32.9% of all variation, and is contributed mainly by median branch length (32.0%), number of nodes (26.2%), height (24.5%), and terminal segment length IQR (11.1%). PC2 captures 21.8% of all variation. Branch length IQR (38.3%), median terminal segment length (23.4%), and circumference (17.9%) are the main contributors to PC2. PC3 captured 20.7% of all variation. Its main contributors are circumference (29.8%), number of branches (29.3%), terminal segment length IQR (17.3%), and median terminal segment length (13.8%).

## Discussion

Our study aimed to examine the ocotillo, a unique desert shrub, in its environmental and ecological context. We did so by modeling and determining the effect of topography and interspecific factors on *F. splendens* morphology and distribution. Past research has attempted to describe the causes and effects of the growth form of ocotillos, but did not test how potential competitors or topography relate to ocotillo morphology. Our approach integrated field data with existing theories of competition dynamics and desert landscape structure, which led to the discovery of novel morphological patterns in the ocotillo. These patterns are not only consistent with these theories, but they are also consistent across our analyses. By examining the ocotillo in this multi-species community and diverse landscape, we collected a wide range of field data and generated novel models to describe *F. splendens* morphology among its nearby neighbors and within its local habitat.

In *F. splendens*, growth occurs either through the generation of new segments, the lengthening of extant segments, or by the production of new branches (Darrow, 1943; Killingbeck, 2016). This relationship was reinforced in PCA1 because the main contributions to PC2 were circumference and number of branches. Conversely, PC1 is contributed mainly by number of nodes, median branch length, and height. This suggests that the lengthening of branches and production of new branches are unrelated. Moreover, from our regressions, we observed that there are multiple morphological features which grow together in an ocotillo; this includes 1) growth between circumference and number of branches and 2) between height, the number of nodes, and median branch length. Ocotillos that grew more branches had larger circumferences. On the other hand, as ocotillos grew more segments, their branches grew in length, and as median branch length increased, ocotillos grew taller. There was also a strong positive relationship between median branch length and terminal segment length IQR, which suggests that recent segment length becomes more variable as ocotillos grow. The effect of age on ocotillo growth form has been recently studied, demonstrating that segment growth becomes more stochastic as ocotillos age (Darrow, 1943; Killingbeck, 2016). Because all ocotillos in this study were either below 2 m in height or had fewer than 40 branches, they would not be mature (Darrow, 1943), but ocotillos growing to maturity could affect intraplant variability as well. Our study instead found that intraplant variability was mediated by site where, for example, only ocotillos along the upper bajada exhibited lesser node counts if they had more variable branch lengths. It is possible that productivity gradients like a bajada, where there is greater water availability at the top of the bajada that decreases with descending elevation (Philips and MacMahon, 1978), would have a stronger influence on ocotillo morphology variability than age.

We also observed how accounting for variables that summarize ocotillo variation can drive intraplant morphological relationships. When IQR variables are added to the PCA analysis as shown in PCA2, the growth relationships between circumference and number of branches, as well as between height, node count, and median branch length are preserved; however, these relationships become less correlated with their PCs, and so they tend to vary less together. The impact of IQR variables on the PCA indicates that the range of branch lengths and terminal segment lengths varies greatly between individuals. However, capturing this variation without masking other effects is one of the main problems surrounding ocotillo morphology (Bowers, 2005; Killingbeck, 2019). For instance, when testing terminal segment length, we found that ocotillos along the upper and lower, plain-like sections of the bajada had conflicting relationships with node growth and segment growth. On the bajada, ocotillo segment length grew with node count while across the bajada and plain the relationship was the inverse. This could possibly be explained by the observation of a wider range of branch length variation across both sites, but when only one site was considered, branch length IQR could be minimized and its effect could fail to override other morphological or interspecific effects. In turn, it appears that localizing ocotillo growth is important in order to observe either weaker or local effects on recent ocotillo growth.

In addition to understanding ocotillo intraplant variability, the lack of relation between height and the number of branches may provide an explanation as to why ocotillos grow intermittently. One possible explanation for this lack of relation could be the association of a potential trade-off where there is a choice between branch production and branch elongation; this choice could depend on the maximization of reproductive output. Stem growth is highly variable, but the production of flowers per stem is relatively consistent (Killingbeck, 2019). Because longer branches produce more flowers (Bowers, 2006), it may not always be advantageous to maximize branch number. It may be advantageous, under certain conditions, to lengthen extant branches to produce more flowers. However, branches can only grow so high due to physical limitations (Ryan and Yoder, 1997); in those cases, growing a new branch would be more advantageous for reproduction. Then, the choice between the production and elongation of branches would depend on maximizing reproductive output; however, further studies would need to quantify ocotillo flower production to ocotillo growth form by, for instance, counting inflorescences.

We also observed a consistent pattern where intraspecific distances and interspecific plant groups affected ocotillo morphology, specifically the number of branches and the terminal segment lengths. We found that ocotillos with major shrubs nearby had fewer branches, but had longer terminal segment lengths. Additionally, the more small cacti surrounding the base of an ocotillo, the shorter the segments were for ocotillos along the upper bajada. These results suggested to us that there is a potential selective pressure, possibly due to competition, that an ocotillo undergoes in order to either produce or lengthen its branches. Local interspecific competition could be mediating this trade-off; a trait theory posited by Ge et al. (2019) explains that trait trade-offs are an indication of organization in an ecological community. In this study, the growth pattern between ocotillo branch number and segment length could reveal community processes like competition. Congruently, we found that ocotillo size influenced the spatial distribution of other ocotillo neighbors. Larger ocotillos with larger circumferences and taller heights had neighboring ocotillos that were farther away, following Pielou’s model (Pielou, 1962), which demonstrates a positive correlation between nearest-neighbor distance and plant size. It is then possible that there is a relationship between competition and ocotillo plant density as suggested by Kadmon (1995), who found a positive correlation between competition and plant density in the desert. However, our study did not determinately reach this conclusion, because the direction of causality between ocotillo size and ocotillo distribution is inconclusive. More isolated ocotillos may grow larger due to higher water availability alongside the arroyo or near the top of the bajada; however, it is also possible that ocotillos curb the growth of other ocotillos as they compete for resources beyond water such as pollinators, root space, or nurse plants for seedling establishment. It has been observed that young ocotillos often had shade protection provided either by an interspecific nurse plant or rock (Nobel and Zutta, 2004). In order to increase its chances to survive and reach maturity, an ocotillo would need to disperse farther from its own kind and establish alongside interspecific nurse plants.

Site, elevation, and distance to the nearest arroyo also influenced ocotillo morphology and distribution. Ocotillos that grew on the plain were on average taller than their upper bajada counterparts. In addition, ocotillos along the bajada were closer to one another and had fewer nodes as elevation increased. In turn, intraspecific distances were modeled most strongly by limited spacing in higher elevation, leading to ocotillos with smaller plant forms. Finally, we observed that distance to the nearest arroyo affected interspecific distances and the number of branches on an ocotillo. The closer an interspecific neighbor was to an arroyo, the further the neighbor was from the nearest ocotillo. As the distance from an ocotillo to an arroyo decreased, the number of branches on the ocotillo increased as well. Thus, these relationships indicate that the factors that relate to ocotillo branch number or segment length are not solely dependent on the limited resources characteristic of desert communities, such as water availability. It appears to be a mix of local factors, including elevation, spatial availability, the proximity of neighbors, and the presence of particular desert plants like shrubs or cacti.

However, there are limitations to our study. The very underlying mechanism for this posited trait trade-off is unknown and our models do not help in elucidating how interspecific spacing occurs. For instance, we found that major interspecific distance was negatively related to distance to the nearest arroyo, suggesting that overall interspecific spacing was mainly mediated by limited resources, but no water availability measurements were taken. Our cross-reference with the hydrographic data from the NDH would not be enough to fully explain desert interspecific neighbor spacing. Similarly, although ocotillos higher on the bajada had closer neighboring ocotillos, leading to more aggregation under limited space, the causality of this aggregation is still uncertain. Others have suggested it could be a mix of root morphology and increasing soil particle size as elevation increases (Yeaton & Cody, 1976). Because the root system of an ocotillo is shallow, reaching a depth of only 7.5-15 cm (USDA Index of Species), it could be in competition with species with deep taproots as previously observed (Cody, 1985). With limited spacing in the upper bajada to spread roots, there could be greater chances of competition among ocotillos and their nearby cacti neighbors.

Age and high intra-plant variability, as referenced before, also continued to be problematic factors while analyzing our data, especially in regards to terminal segment length. This is not a new problem, as it has been shown that variation among branches on a single ocotillo overwhelmed any differences between individual ocotillos (Bowers, 2005). Our mixed effect models helped explain relationships between segment length and interspecific neighbor presence or count, but they had conflicting relationships across different sites. Nonparametric tests and nonparametric factors such as the terminal segment length IQR and median branch length IQR made it possible to observe those relationships. However, future research will need to use factors that decrease variation within the ocotillo but increase interpretive and descriptive power in order to better understand ocotillo morphology and growth. If larger sample sizes were used and more ocotillos on topographically different sites were measured, we would expect our principal components to better reflect our linear models.

Ultimately, the indication of a trade-off between branch number and terminal segment length, and their modeled relationships between ocotillo morphology, interspecific and intraspecific distances, and topography opens paths to further explore their causal effects. Our trade-off indication may help explain previous observations on the irregularity of *F. splendens* terminal segment formation (Killingbeck, 2017) by revealing novel factors impacting ocotillo morphology. However, we encourage further testing of ocotillo growth patterns along various topographically different sites. In addition, tests on the fitness and genetic basis of ocotillo growth are clearly needed to definitively assess our interpretations. Accurate predictions of ocotillo growth and spacing will likely rely on models that incorporate both individual traits and extrinsic environmental factors. Overall, our approach and results can better help frame desert species within their environmental and ecosystem dynamics and can be used to better understand the role and impact of ocotillos in desert communities.

We would like to thank Colin Kyle and Maia McNeil for assistance with field morphology measurements. We would like to thank the Organ Pipe Cactus National Monument for permission to study *Fouquieria splendens* within its national monument.

---

Supplementary Table 1--Multivariate model and mixed effect models of *Fouquieria splendens* morphology for ocotillos located on a bajada and a plain in Organ Pipe National Monument, Arizona. All models were grouped by their response variable and ordered by their ascending AIC values. In the “Dataset” column, “all” indicates a mix of bajada and plain ocotillos while “bajada” indicates only ocotillos measured on the bajada. Ocotillos located on the bajada were encoded with site = 0 while ocotillos on the plain were encoded with site = 1. Interspecific plants are split between two groups – shrub and cactus – where cactus = 0 and shrub = 1.

Supplementary Table 2--Best fit models of *Fouquieria splendens* morphology for ocotillos located on both a bajada and a plain in Organ Pipe National Monument, Arizona. In the “Dataset” column, “all” indicates a mix of bajada and plain ocotillos while “bajada” indicates only ocotillos measured on the bajada. Significant predictor variables (*P* < 0.05) are listed.

Supplementary Table 3--Multivariate models of interspecific and intraspecific distance to *Fouquieria splendens* located on both a bajada and a plain in Organ Pipe National Monument, Arizona. All models were grouped by their response variable and ordered by their ascending AIC values. In the “Dataset” column, “all” indicates a mix of bajada and plain ocotillos while “bajada” indicates only ocotillos measured on the bajada. Interspecific plants are split between two groups – shrub and cactus – where cactus = 0 and shrub = 1.

Supplementary Table 4--Best fit models of interspecific and intraspecific distance to *Fouquieria splendens* located on both a bajada and a plain in Organ Pipe National Monument, Arizona. In the “Dataset” column, “all” indicates a mix of bajada and plain ocotillos while “bajada” indicates only ocotillos measured on the bajada. Significant predictor variables (*P* < 0.05) are listed.

## Literature Cited

Abdi, H., and Williams, L. 2010. Principal component analysis. Wiley Interdisciplinary Reviews: Computational Statistics 2:433–459.

Bowers, J. E., and McLaughlin, S. P. 1982. Plant species diversity in Arizona. Madroño 29:227–233.

Bowers, J. E. 2005. Influence of plant size and climatic variability on the floral biology of Fouquieria splendens (ocotillo). Madroño 52:158–165.

Bowers, J. E. 2006. Branch length mediates flower production and inflorescence architecture of Fouquieria splendens (ocotillo). Plant Ecology 186:87–95.

Bowers, M. A. and Lowe, C. H. 1986. Plant-form gradients on Sonoran Desert bajadas. Oikos 46:284–291.

Búrquez, A. et al. 1999. Vegetation and habitat diversity at the southern edge of the Sonoran Desert. The University of Arizona Press. 36–67.

Darrow, A. R. 1943. Vegetative and floral growth of Fouquieria splendens. Ecology 24:310–322.

Fonteyn, P. J. and Mahall, B. E. 1978. Competition among desert perennials. Nature 275:544–545.

Fowler, N. 1986. The role of competition in plant communities in arid and semiarid regions. Annual Review of Ecology, and Systematics 17:89–110.

Gaudet, C., and Keddy, P. A. 1989. Plant Competition. Nature 337:123.

Ge X. M., et al. 2019. Functional trait-trade-off and species abundance: insights from a multi-decadal study. Ecology Letters. 1–10.

Grime, J. P. 1973. Competitive exclusion in herbaceous vegetation. Nature 242:344–347.

Huston, M. A. 1979. A general hypothesis of species diversity. American Naturalist 113:81–1001.

Ji, W., et al. 2019. Constraints on shrub cover and shrub-shrub competition in a U.S. southwest desert. Ecosphere 10:1–16.

Kaiser, H. E. 1960. The application of electronic computers to factor analysis. Education & Psychological Measurement 20: 141–151.

Kaufman, J. D. and Dunlap, W.P. 2000. Determining the number of factors to retain: A windows-based FORTRAN-IMSL program for parallel analysis. Behavior Research Methods, Instruments, & Computers 32:389–395.

Kadmon, R. 1995. Plant competition along soil moisture gradients: A field experiment with the desert annual Stipa capensis. Journal of Ecology 83:253–262.

Killingbeck, T. K. 2006. Loss of long-shoot leaves may mask terminal stem segment age in ocotillo (Fouquieria splendens). BioOne Complete 12:11–12.

Killingbeck, T. K. 2009. Flowers are produced on stems shorter than one meter in the desert shrub ocotillo (Fouquieria splendens). The Southwestern Naturalist 54:55–57.

Killingbeck, T. K. 2016. Tracking growth and age in the drought-deciduous shrub Fouquieria splendens (ocotillo) in the Chihuahuan Desert: The role of wood rings and stem segments. The Southwestern Naturalist 61:217–224.

Killingbeck, T. K. 2017. Are growth rings accurate fingerprints of plant age in a stem-succulent, drought-deciduous shrub growing in the Chihuahuan Desert? Journal of Arid Environments 144:220–223.

Killingbeck, T. K. 2019. Stem succulence controls flower and fruit production but not stem growth in the desert shrub ocotillo (Fouquieria splendens). American Journal of Botany 106:223–230.

Lundgren, M. R., and Marais, D. D. 2020. Life history variation as a model for understanding trade-offs in plant-environment interactions. Current Biology 30:180–189.

McAuliffe, J. R. 1994. Landscape evolution, soil formation, and ecological patterns and processes in Sonoran Desert bajadas. Ecological Monographs 64:111–148.

Nobel, P. S. and Zutta, B. R. 2004. Morphology, ecophysiology, and seedling establishment for Fouquieria splendens in the northwestern Sonoran Desert. Journal of Arid Environments 62:251–265.

Nobel, P. S. and Zutta, B. R. 2007. Rock associations, root depth, and temperature tolerances for the “rock live-forever,” Dudleya saxosa, at three elevations in the north-western Sonoran Desert. Journal of Arid Environments 69:15–28.

Pearse, I.S, et al. 2020. Life-history plasticity and water-use trade-offs associated with drought resistance in a clade of california jewelflowers. The American Naturalist 195:691–704.

Philips, D. L. and MacMahon, J. A. 1978. Gradient analysis of a Sonoran Desert bajada. The Southwestern Naturalist 23:669–679.

Philips, D. L. and MacMahon, J. A. 1981. Competition and spacing patterns in desert shrubs. Journal of Ecology 69:97–115.

Pielou, E. C. 1962. Runs of one species with respect to another in transects through plant populations. Biometrics 18:579–593.

Pierce, N. 2019. Competition suppresses shrubs during early, but not late, stages of arid grassland-shrubland state transition. Functional Ecology 33:14800–1490.

Ryan, M. G. and Yoder, B. J. 1997. Hydraulic limits to tree height and tree growth. BioScience 47:235–242.

Stromberg, J. C. 2007. Seasonal reversals of upland-riparian diversity gradients in the Sonoran Desert. Diversity and Distributions 13:70–83.

